# Recellularization of bronchial extracellular matrix with primary bronchial smooth muscle cells

**DOI:** 10.1101/2019.12.23.878645

**Authors:** Selma Ben Hamouda, Amandine Vargas, Roxane Boivin, Maria Angelica Miglino, Renata Kelly da Palma, Jean-Pierre Lavoie

**Author notes:** Corresponding authors (JPL) (SBH).

## Abstract

Severe asthma is associated with an increased airway smooth muscle (ASM) mass and an altered composition of the extracellular matrix (ECM). Studies have indicated that ECM-ASM cell interactions contribute to this remodeling and its limited reversibility with current therapy. Three-dimensional matrices allow the study of complex cellular responses to different stimuli in an almost natural environment. Our goal was to obtain acellular bronchial matrices and then develop a recellularization protocol with ASM cells. We studied equine bronchi as horses spontaneously develop a human asthma-like disease. The bronchi were decellularized using Triton/Sodium Deoxycholate. The obtained scaffolds retained their anatomical and histological properties. Using immunohistochemistry and a semi-quantitative score to compare native bronchi to scaffolds revealed no significant variation for matrixial proteins. A DNA quantification and electrophoresis indicated that most of DNA was 29.6 ng/mg of tissue ± 5.6 with remaining fragments of less than 100 bp. Primary ASM cells were seeded on the scaffolds. Histological analysis after recellularization showed that ASM cells migrated and proliferated primarily in the decellularized smooth muscle matrix, suggesting a chemotactic effect of the scaffolds. This is the first report of primary ASM cells preferentially repopulating the smooth muscle matrix layer in bronchial matrices. This protocol is now being used to study the molecular interactions occurring between the asthmatic ECMs and ASM to identify effectors of asthmatic bronchial remodeling.

## Introduction

Asthma is a progressive and multi-component respiratory syndrome. Remodeling of the airways in asthma is characterized by structural changes leading to a thickening of the bronchial wall, airflow obstruction, and hyperreactivity of the airways [1]. In human asthma, tissue remodeling is believed to be only partially reversible or even irreversible following conventional treatments (corticosteroids / bronchodilators) even during extended periods of remission [2]. While airway remodeling is considered a target for asthma, little is known of the mechanisms involved in its development and reversibility. This is due to ethical considerations related to the invasiveness when sampling the airways, and technical limitations of the current imaging techniques.

As compared to single-layer culture, three-dimensional (3D) cell culture improves different cellular parameters including viability, adhesion, proliferation, etc. A study comparing the culture of skin fibroblasts on natural 3D matrices to their monolayer culture on different substrates revealed significant variations. Cell adhesion was 5 to 6 times greater on 3D matrix than on single-layer culture, cell migration was also increased and the acquisition of a morphology mimicking the in vivo appearance of the cells occurred faster [3]. A study of human embryonic stem cell differentiation found that cells were physiologically and morphologically more representative of the native cells when cultured on a 3D matrix than when cultured on flasks [4]. This is also supported by the finding that pulmonary fibroblasts response to tumor necrosis factor (TNF)α was increased in 3D culture when compared to monolayer culture [5]. There are different types of 3D culture including: 1) in suspension, 2) in gel scaffold or, 3) in natural or synthetic fibrous scaffold [6, 7]. Culturing cells on 3D bases is more difficult and requires longer culture time than in monolayer, but the results obtained are believed to be more representative of the natural environment [6]. Thus, in order to evaluate the behavior of airway smooth muscle (ASM) cells in the asthmatic airways, we aimed to decellularize an equine respiratory bronchus while maintaining its architecture and protein composition to allow recellularization by bronchial smooth muscle cells. Equine bronchi were studied as horses spontaneously develop an asthma-like condition that shares clinical and remodeling features with human asthma [8]. Furthermore, it was shown in this model that the quantity of ASM is increased, and only partially reversible even after 1 year of inhaled corticosteroids [9, 10]. Results of the present study suggest that the bronchial smooth muscle cells preferentially colonize the bronchial smooth muscle extracellular matrix. These findings may allow investigating the interactions between smooth muscle cells and the extracellular matrix in the asthmatic airways.

## Material and methods

### Animals

Archived lung tissues from four asthmatic and three healthy control horses (5 mares and 2 geldings) aged 10-12 years from a tissue bank (http://www.btre.com) were studied. Four additional lungs were obtained from a slaughterhouse. The experimental protocol was approved by the ethical committee of the University of Montreal number Rech-1578.

### Bronchi decellularization

Bronchi from 2^nd^ to 4^th^ generation were dissected from the surrounding lung tissues within 2 hours after euthanasia. The bronchi were then snap frozen in liquid nitrogen and kept at −80°C until used. These bronchi were thaw and decellularized using a protocol previously described [11] with minor modifications. Briefly, two consecutive cycles of detergent (Triton 1X, Sodium deoxycholate and sodium chloride) and enzymatic (DNase) treatments were followed by sterilization with paracetic acid-ethanol under continued agitation to ensure the elimination of any immunogenic cellular material that may hinder recellularization. Sections of bronchi were then paraffin-embedded for histology or snap frozen for DNA and protein isolation.

### Airway smooth muscle cell isolation and culture

Airway smooth muscle (ASM) cells were isolated from the same horses in the first hour after the death, as previously described [12]. In brief, the ASM layer was collected from the first bronchial bifurcation and then immersed in a digestion medium (Dulbeco’s Modified Eagle Medium / F12 nutrient mix (Thermofisher, Waltham, MA) with 0.125 U/ml Collagenase H (Sigma Aldrich, St. Louis, MO), 1 mg/ml Trypsin inhibitor (Sigma Aldrich, St. Louis, MO), 1 U/ml elastase (Worthington biochemical, Lakewood, NJ), 1% Penicillin-Streptomycin (Wisent Inc., Saint-Jean-Baptiste, QC) and 0.1% Fungizone (Fisher Scientific, Hampton, NH)).

Cells (ASM) were seeded into ventilated cell culture flasks at 300,000 cells/cm^2^ in DMEM/F12 medium supplemented with 0.0024 mg/ml adenine, 10% non-decomplemented fetal bovine serum (FBS) (Wisent Inc., Saint-Jean-Baptiste, QC), 1% Penicillin-Streptomycin and 0.1% Fungizone and cultured at 37°C and 5% CO_2_ for 48 hours. Media was then changed every 48 hours until confluence was reached. Cells were frozen between the first and 4^th^ passage (P) in liquid nitrogen until being used.

### Smooth muscle cells characterization

ASM cells were characterized by flow cytometry before recellularization, as previously described [12]. Briefly, cells were stained for intracellular markers with anti-α-SMA (mouse IgG2a, Sigma Aldrich, St. Louis, MO, 1/250), anti-desmin (rabbit polyclonal IgG, Abcam, Cambridge, UK, 1/200) and anti-SMMHC (rabbit IgG, Biomedial technologies, Stoughton, MA, 1/300) antibodies for 1 hour. Cells were then washed 3 times and incubated for 30 minutes in the dark with fluorescent dye-conjugated anti-IgG antibodies. Isotype-matched control antibodies (mouse IgG2a and rabbit IgG) were used as negative control. All signals greater than those of the isotype-matched control antibodies were considered positive, and degree of staining was evaluated as the mean fluorescence intensity and mean percentage of positive cells. This characterization showed simultaneous expression of α-SMA (mean±SEM) for 90% ± 8.6 cells, SMMHC for 71% ± 16 cells and desmin for 85 ± 9.2 cells.

### Assessment of decellularization efficiency

Decellularized bronchi were stained and compared to native bronchi using the Russel modification of Movat Pentachrome [13]. The protocol was modified as the exposure time to ferric chloride and to alcoholic safran solution was changed to 1 and 5 minutes, respectively. Images were obtained at 100 and 200 magnifications using Panoptiq software (version 2) connected to a Prosilica GT camera (model: GT1920C) mounted on a Leica DM4000 B microscope.

DNA was isolated from 10 mg of frozen native and from freshly decellularized bronchi using DNeasy blood and tissue Kit^®^ (Invitrogen, Hilden, DE) as recommended by the manufacturer. DNA was then visualized on agarose gel. Quantification of doublestranded DNA before and after decellularization was done using the Qubit DNA BR Assay kit (Invitrogen, Carlsbad, CA) according to the manufacturer instructions. Proteins were extracted using T-PER (Thermofisher, Waltham, MA) and quantified using Qubit Protein Assay Kit (Invitrogen, Carlsbad, CA).

Immunohistochemical staining for collagen I, collagen IV and fibronectin was performed on 10% formalin preserved native and decellularized bronchi. Tissues were incubated overnight with primary antibodies (collagen I; rabbit anti-bovine IgG, Cederlane, Burlington, ON, dilution 1:500, collagen IV; mouse anti-human IgG, Dako, Carpinteria, CA, dilution 1:50, and fibronectin; unconjugated rabbit polyclonal antibody, Biorbyt, San Francisco, CA, dilution 1:150). The biotinylated secondary antibodies were applied at the same concentrations as the primary antibodies for 45 minutes. Vectastain ABC kit (Biolynx, Brockville, ON) was applied before DAB revelation (Vector Laboratories, Peterborough, UK) and a counterstain with Harry’s hematoxylin. Negative controls were also prepared. They were stained with rabbit or mouse IgG instead of the primary antibodies to reveal potential unspecific staining. Using the negative controls as a benchmark, a semi-quantitative score was established for the basement membrane, smooth muscle, blood vessel and lamina propria labeling as follows: Grade 0: Absence of staining, Grade 1: Presence of staining. From this score, an average was established to compare the labeling difference between native and decellularized bronchi.

### Bronchi recellularization protocol

Decellularized bronchi were split in two and secured on a sterile support, then cut into small pieces of a maximum of 1×1cm and rinsed in sterile PBS 1X. Tissues were then placed in a 24-well plate (Costar, Washington, D.C.) and recellularized with the ASM cells between P4 and P7 at a concentration of 158,000 cells/cm^2^. One and a half milliliters of medium were added to the culture under the same condition described above. After a 48-hour incubation, allowing primary adhesion, 1 ml of the medium was changed in each well. Then, the medium was changed every other day. Tissues were maintained in culture, in the same well, between 48 hours and 41 days or transferred at 31 days to a 6-well plate (Celltreat, Pepperell, MA) for 10 more days. Samples were collected at day 2, 7, 14, 21 and 31 without tissue transfer and at day 41 with and without tissue transfer (Supplemental figure 1).

### Assessment of recellularization efficiency

Tissues were fixed in 10% formalin, paraffin-embedded then sliced at 4.5 μm thickness and stained using Movat Pentachrome histological staining protocol. The qualitative assessment of recellularization was based on a visual examination of the recellularized tissue sections under the optical microscope at 100 and 200 magnifications using the Panoptiq software, as previously described.

An immunofluorescence staining of 5 fresh-frozen bronchi recellularized at day 41 was performed for α-SMA. Tissues were incubated with the primary antibody for 2 hours at 37°C (α-SMA anti-mouse IgG2a, 1:250, Sigma Aldrich, St-Louis, MO). The fluorescent secondary antibody (Goat anti-mouse IgG, 1:1000, Invitrogen, Carlsbad, CA) was incubated for 1 hour at room temperature. An isotype control was used as a benchmark for positive staining. The slides were analyzed under Olympus Fluoview FV1000 confocal unit attached to the inverted Olympus IX81 microscope (Olympus Canada, Richmond Hill, ON, Canada) and compared to their replicate on a Movat Pentachrome staining to confirm the histological results.

For scan electron microscopy, the recellularized bronchi were fixed in 2.5% glutaraldehyde, washed and post-fixed in 1% aqueous osmium tetroxide solution, and dehydrated in increasing series of alcohol (70% to 100%). After dehydration, the samples were dried on the LEICA EM CPD 300 Critical Point Apparatus, mounted on carbon tape and gold-plated on the Emitech K550 Metalizing Apparatus, photo-documented on the LEO 435VP Scanning Electron Microscope at the Advanced Diagnostic Center by Image - CADI - Faculty of Veterinary Medicine and Animal Science - University of São Paulo.

### Statistical analyses

The values are expressed as mean ± standard error of the means (SEM). Values of DNA and total protein quantification were analyzed by use of the paired student test (GraphPad Prism 7). Immunohistochemistry scores were analyzed by an exact chi-square test to compare the prevalence of positive staining against the status of the bronchi (native or decellularized) using SAS v.9.3. Values of P ≤ 0.05 were considered significant.

## Results

### Decellularization efficiency assessment

#### Histological assessment

Visual examination under the optical microscope at magnifications 100 and 200 confirmed the absence of cellular structures in decellularized matrices in comparison to the native bronchi (Fig 1). The epithelial cell layer was totally removed. It also showed a preserved bronchial architecture after the decellularization process with a maintenance of the tissue organization and the contents in collagen and elastic fibers.

**Fig 1.**
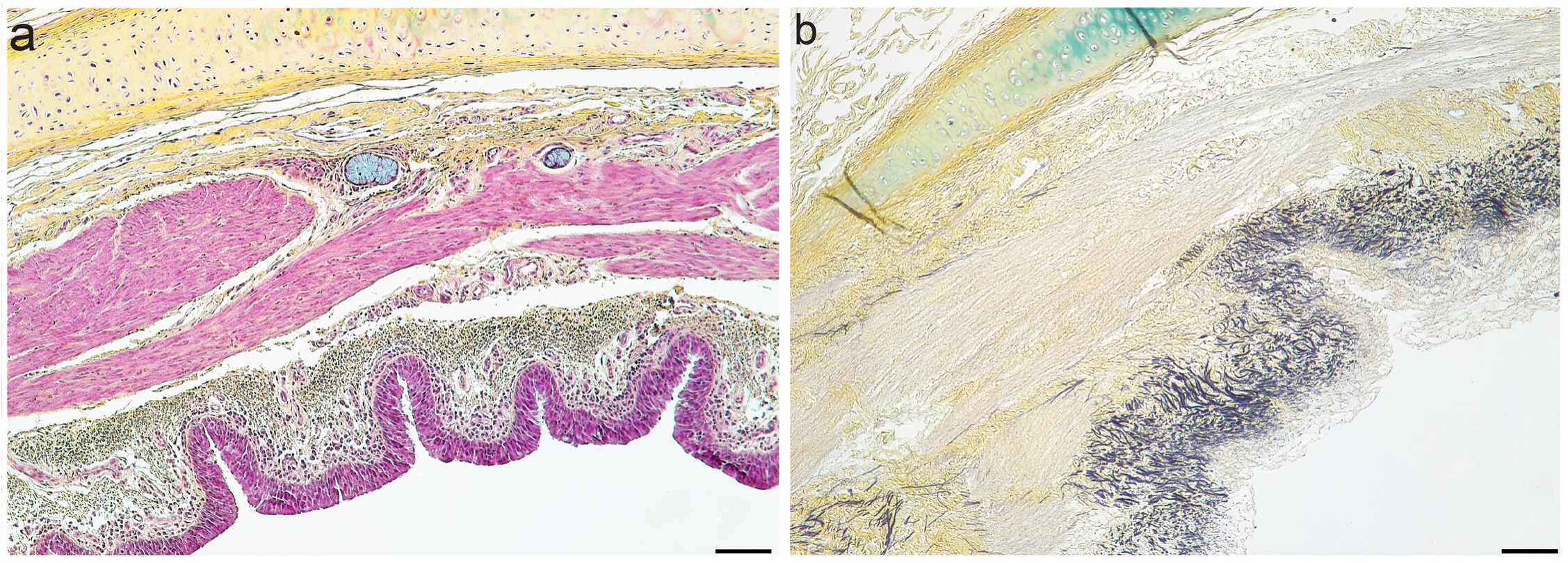
Movat Pentachrome staining of a native and a decellularized bronchus at magnification 100. (a) native bronchus, (b) decellularized bronchus. From top to bottom, on each image, the bronchial cartilage, followed by the bronchial smooth muscle surrounded on both sides by the extracellular matrix which, at the level of the lamina propria, is followed by the epithelial cells layer. Collagen is stained in yellow. The black filamentous structures on (a) and on (b) represent the elastic fibers. The cell nuclei are stained in purple, their omnipresence is noted on (a) and their total absence on (b). Scale bars indicate 100 μm.

#### DNA quantification and electrophoresis

A decrease in DNA concentration was observed in decellularized bronchi. The mean DNA concentration in the native bronchi was 2529 ng/mg of tissue ± 72.7 whereas it was 29.6 ng/mg of tissue ± 5.6 (p < 0.0001; Fig 2a) after decellularization. Agarose gel electrophoresis revealed that the remaining double-stranded DNA fragment lengths in decellularized bronchi was less than 100 bp (Supplemental figure 2).

**Fig 2.**
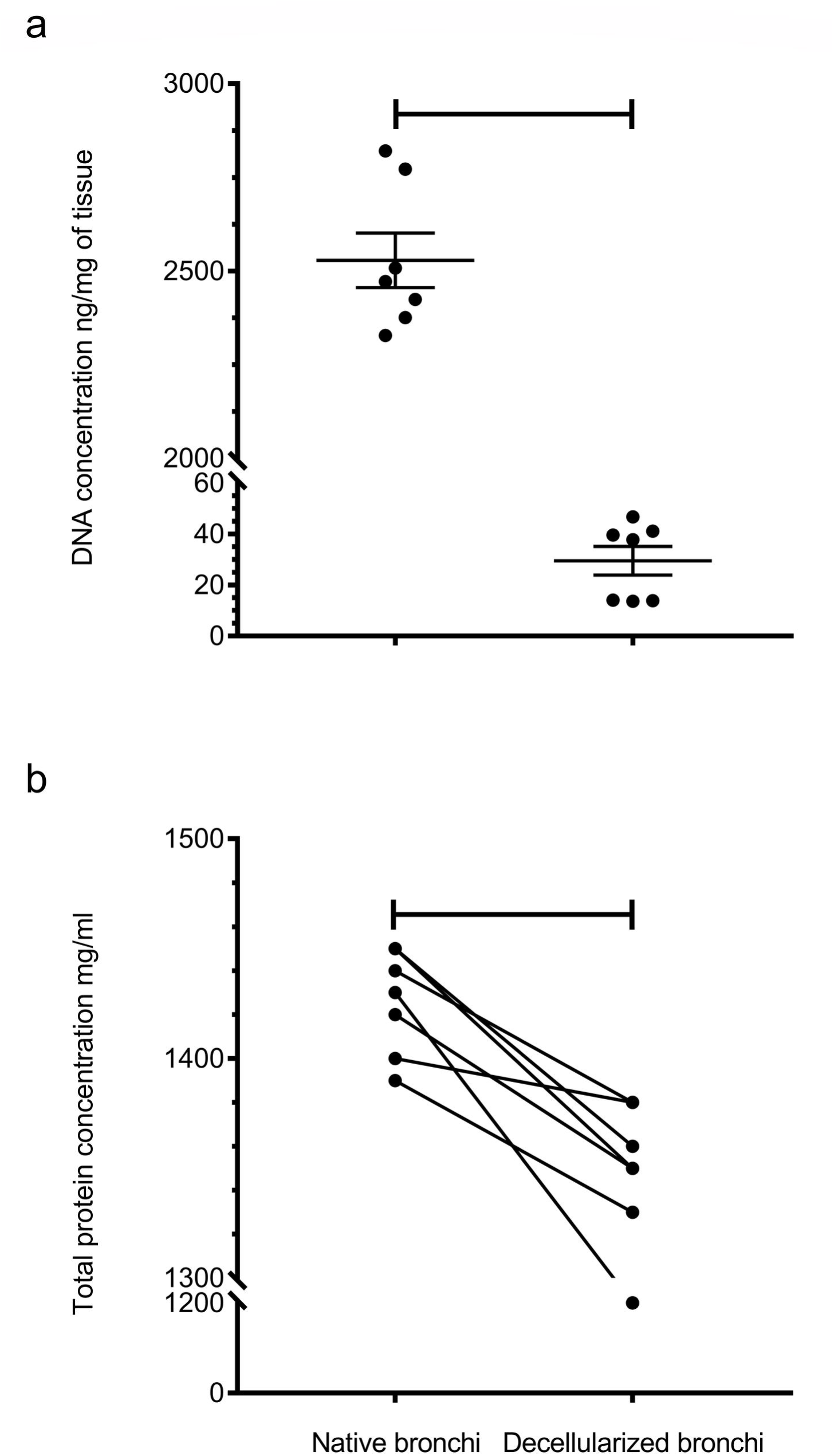
DNA and total protein concentrations in native and decellularizd bronchi. (a) DNA concentration in ng/mg of tissue in 7 native and decellularized bronchi, (b) Total protein concentration in mg/ml in 7 native and decellularized bronchi. A significant difference is found for DNA and total protein concentrations between native and decellularized bronchi with respectively, p < 0.0001 and p = 0.01.

#### Protein quantification

Although the difference in total protein concentrations in the native and decellularized bronchi was significant (p = 0.01), the decline in these concentrations remained moderate, to approximately 100 mg/ml (Fig 2b). The qualitative and semi-quantitative evaluation revealed that collagen I and IV are not affected by the decellularization process. The fibronectin while decreased, remained detectable in abundant amounts in the decellularized extracellular matrix (Fig 3k–3l).

**Fig 3.**
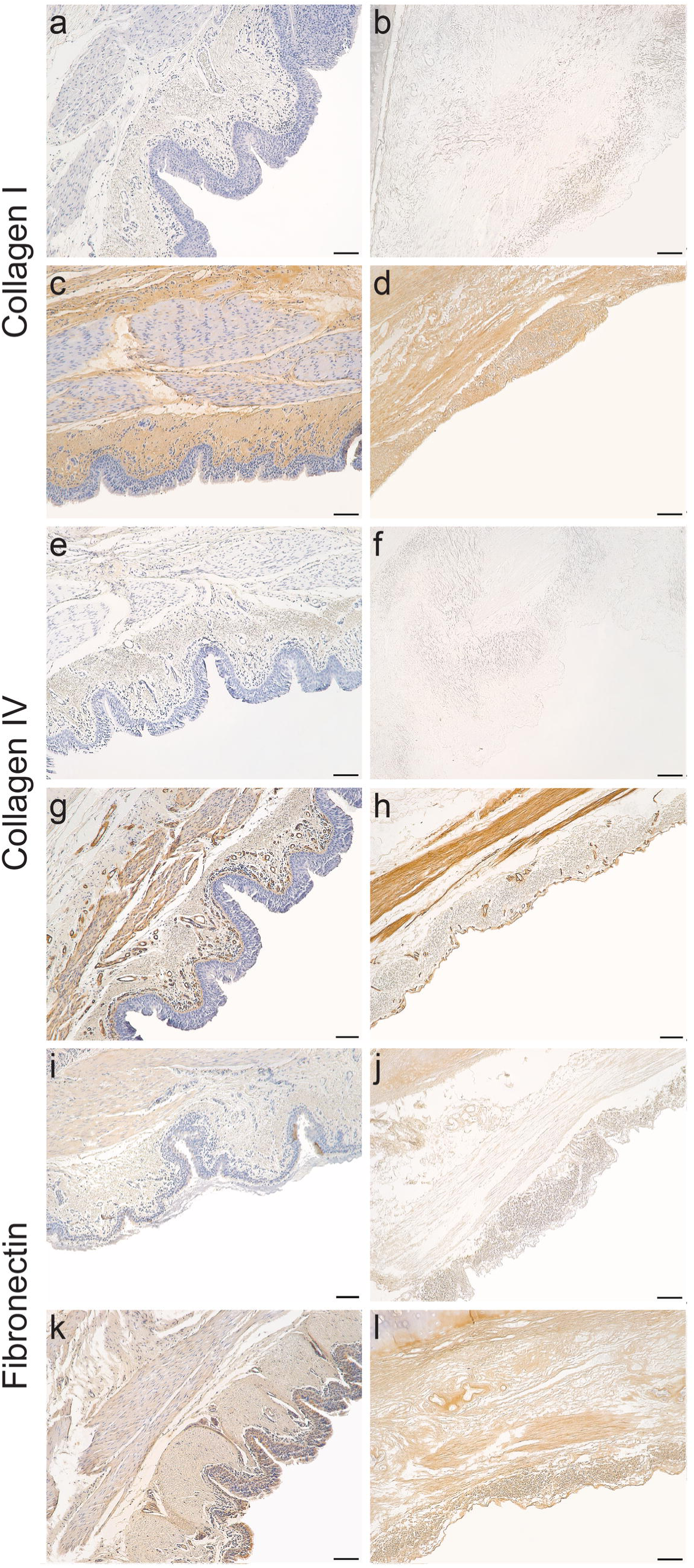
Immunohistochemical staining for Collagen I and IV and for fibronectin in native and decellularized bronchi at magnification 100. Native and decellularized bronchi stained for collagen I (a, b, c, d), collagen IV (e, f, g, h) and fibronectin (i, j, k, l). Evaluation of positive staining (c, d, g, h, k, l) compared to isotypic controls (a, b, e, f, i, j). Scale bars indicate 100 μm.

#### Recellularization assessment

Five different equine bronchi were recellularized with 3 primary ASM cell lines between passages 3 and 7. Recellularizations from 48 hours to 7 days were modest or absent in all cell-tissue combinations tested (n = 19). However, between days 14 and 21, the ASM cells were detectable in tissue in 17 out of 19 replicates made during 7 different recellularization trials.

At 31 days, ASM cells were observed in all recellularized tissues (n = 20). On 4 different recellularization assays, the amount of ASM cells within the scaffold was maximum at 41 days. The ASM cells that repopulated the decellularized bronchi were first located on the surface of the basement membrane or in the extracellular smooth muscle matrix (14 days). At day 21, ASM cells were present in the smooth muscle layer of the decellularized bronchi and appeared to colonize it preferentially (Fig 4). This was confirmed between 31 and 41 days, with abundant cells in the muscular extracellular matrix and adjacent to the bronchial cartilage. However, some cells remained located above the basement membrane. In two recellularization trials with 3 biological replicates, the recellularization was improved at 41 days by transferring the tissue at day 31 in another well for 10 additional days of culture before harvesting.

**Fig 4.**
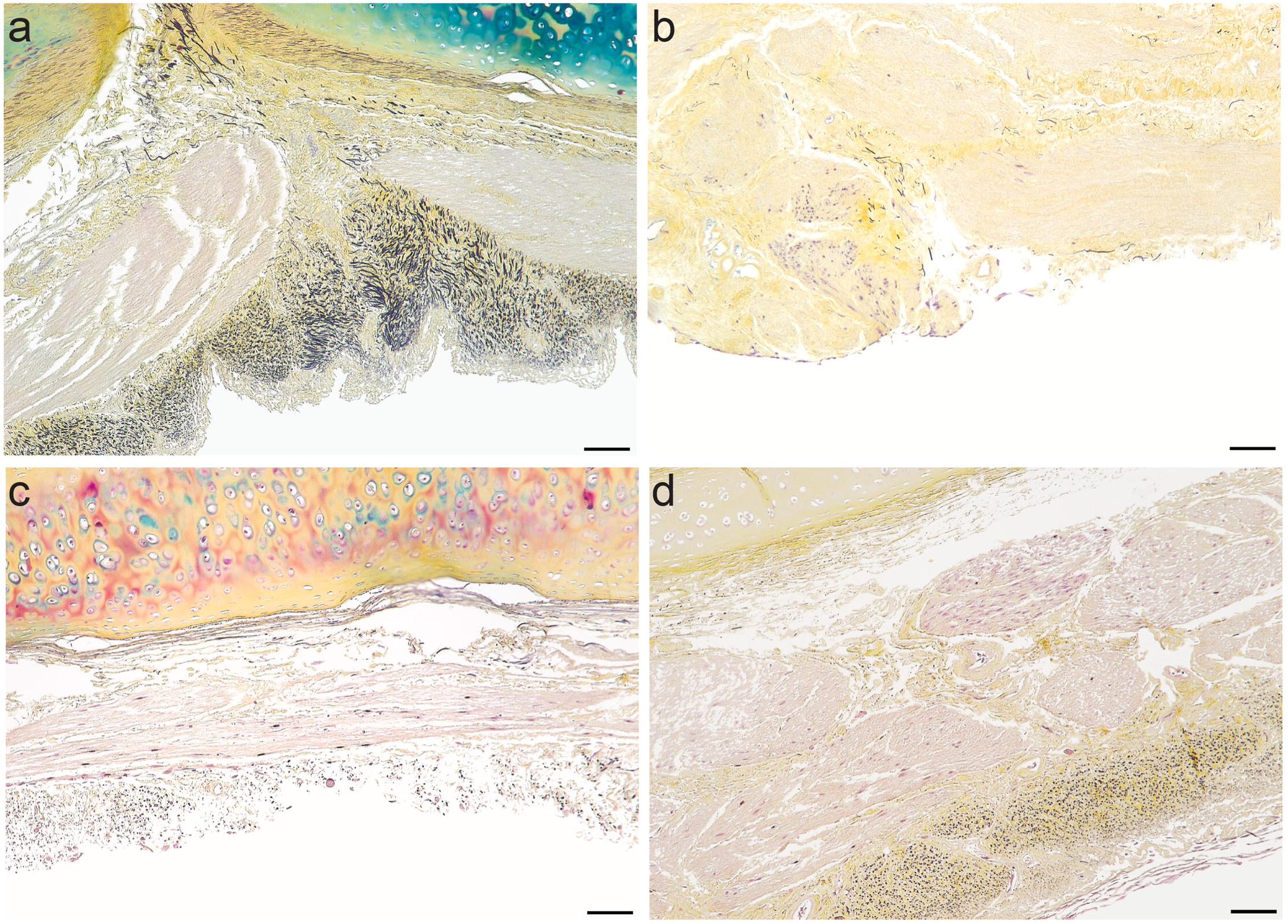
Movat Pentachrome staining at magnification 100 showing the recellularization between day 0 and day 41. The ASM cell nuclei are stained in purple. (a) decellularized bronchi on day 0; (b) recellularized bronchi on day 14 with ASM cells at the surface and in the collagenous ECM of the smooth muscle; (c) recellularized bronchi on day 21 with ASM cells more organized at the level of the smooth muscle ECM and lined with bronchial cartilage; (d) recellularized bronchi on day 41 with confluent ASM cells in the smooth muscle extracellular matrix and on the surface of the basement membrane. Scale bars indicate 100 μm.

The immunofluorescent staining confirmed the expression of α-SMA by the cells present in zones of abundant recellularization in the smooth muscle matrix (Fig 5).

**Fig 5.**
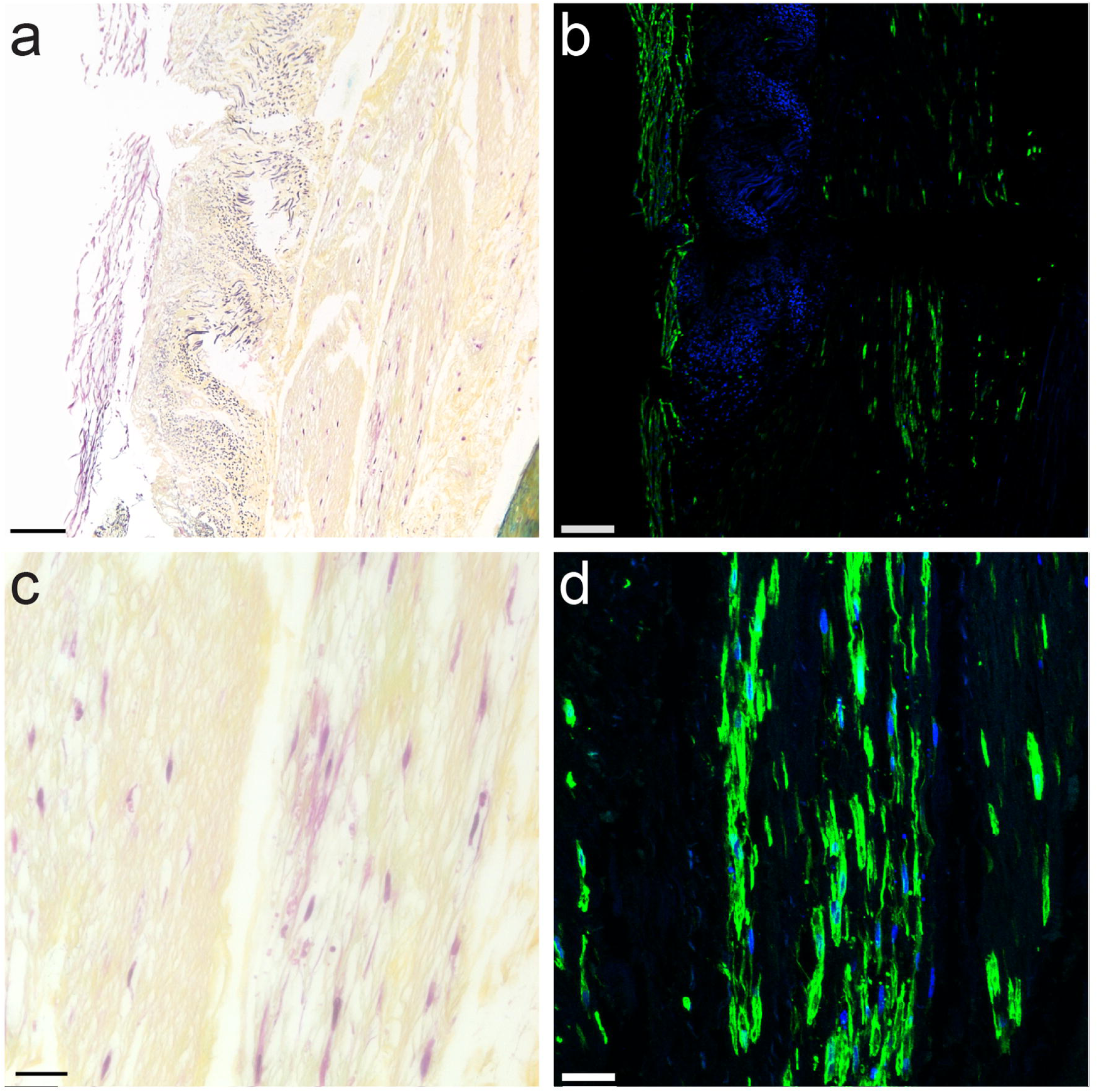
Movat Pentachrome and α-SMA immunofluorescence consecutive staining of recellularized bronchial matrix at day 41. (a) and (b): Movat Pentachrome and immunofluorescence staining of the same area in a recellularized bronchus at magnification 100. (c) and (d): Movat Pentachrome and immunofluorescence staining of the same area in a recellularized bronchus at magnification 400. The cells recellularizing the tissue are expressing the α-SMA which is an indication of the smooth muscle nature of the cells colonizing the tissue and their purity. Scale bars are indicating 100 μm for (a) and (b) and 25 μm for (c) and (d).

On scanning electron microscopy analysis, recellularization was identified in most samples on the tissue surface. On some parts of the ECM, where the resistance seamed reduced, the cells penetrated the tissues, as it is shown, Fig 6.

**Fig 6.**
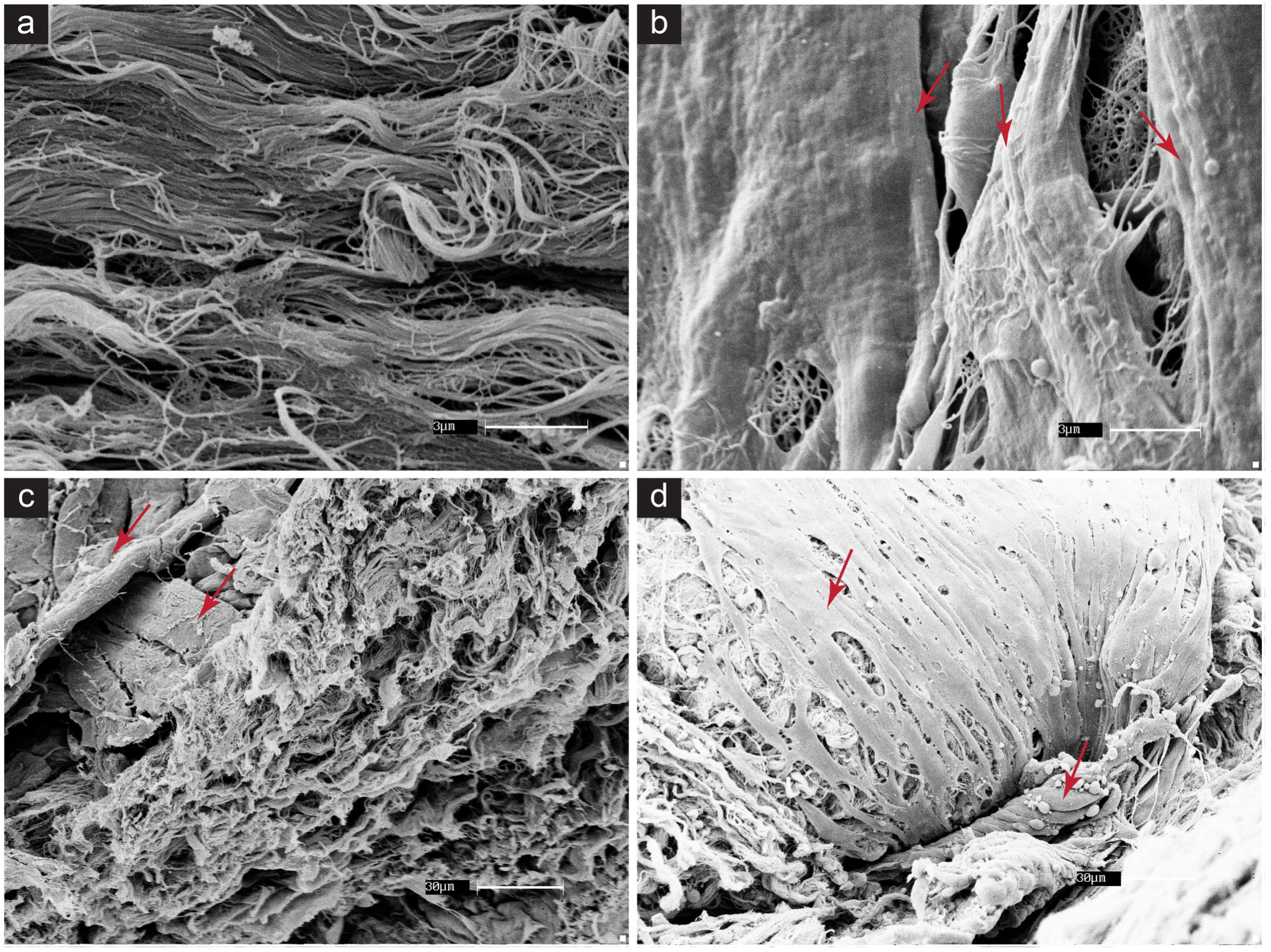
Scan electron microscopy of decellularized and 41 days recellularized bronchial matrices. (a) decellularized matrix; (b) recellularized matrix with cells appearing on the surface; (c) cross section of recellularized matrix, cells are only on the surface but not inside the tissue; (d) cross section of recellularized matrix, cells are penetrating the tissue. The red arrows are indicating some of the cells. Scale bars are indicating 3 μm for (a) and (b) and 30 μm for (c) and (d).

## Discussion

In this study, we developed a protocol to recellularize decellularized respiratory bronchi with ASM cells. The results showed a preferential migration and colonization of the muscular extracellular matrix by these cells. This phenomenon is, to the best of our knowledge, reported for the first time in any species. The protocol presented herein will enable the study of the phenotypic changes of ASM cells by an asthmatic ECM.

The initial aim of tissue engineering was to develop organs devoid of immunogenic rejections for transplantation [14, 15]. It was then adopted in pharmacological and oncological researches to identify the cellular response to drugs and in diseases [16–18], as the cells’ behavior was shown to vary depending on the 3D structures of the substrate [6, 19]. Different organs, including lung tissues, have been decellularized and then recellularized [20–23], mainly with stem cells [24, 25]. Recent studies have recellularized horse lung tissues with equine dermal fibroblasts and canine yolk sac cells [26], and mouse lungs with human and murine fibroblasts [27]. McClure *et al.* grafted a decellularized skeletal muscle in the gastrocnemius and demonstrated a regeneration of the graft with the presence of cells and neuromuscular junctions [28]. However, none of the studies reported a preferential recellularization for specific cell types when seeded on a heterogenous biological matrix.

To the best of knowledge, the present study is the first to attempt to recellularize airways with primary airway smooth muscle cells. Different combinations of smooth muscle cells lineages and matrices were studied and resulted in the colonization of the bronchial smooth muscle matrix by the ASM cells. This preferential colonization likely involved complex cellular mechanisms including integrin expression, adhesion, migration and proliferation and suggests that the decellularized tissue retained enough of its native qualities to allow this process to occur. Da Palma *et al.* demonstrated that the fibroblasts recellularizing a decellularized horse lung are expressing the N-cadherin, an adhesion biomarker [26]. This protein may play an important role in the ASM cell migration seen in our study as it has been shown that the migration of the vascular smooth muscle cells is delayed by the inhibition or the down-regulation of the N-cadherin [29, 30]. Moreover, the ECM is known to regulate the migration of the ASM cells [31, 32].

The smooth muscle colonization was uneven as some parts of the bronchi contained more ASM cells than other. This unevenness in cell distribution has also been observed during lung recellularization by endothelial cells [33]. Uygun *et al.* also described variable hepatocytes distribution within hepatic matrix [34]. This may be due to the uneven mechanical properties of the decellularized bronchi as scan electron microscopy results suggest that cells seem to reach the smooth muscle ECM from zones with low tissue resistance.

The presence of matrikines in the scaffolds may also have contributed to this preferential cell colonization. Matrikines are peptides produced from the proteolytic degradation of the extracellular matrix [35]. Given the cellular destruction that occurs during decellularization, the release of intracellular proteases may have resulted in the production of a high concentration of matrikines within the scaffolds. These peptides would affect cell behavior across integrins by stimulating the secretion of certain growth factors. It has been shown that valine-glycine-proline-valine-glycine (VGPVG), a hydrophobic elastin matrikine, stimulates smooth muscle cell proliferation [36]. The tripeptide sequence Arginine-Phenylalanine-Lysine (RFK) derived from thrombospondin-I is also mitogenic to smooth muscle cells through the activation of the transforming growth factor (TGF)β [37]. Another peptide, valine-glycine-valine-arginine-proline-glycine (VGVAPG), is chemotactic for fibroblasts [38]. These matrix fragments being mobile and regulating cell behavior, would be potential promoters of myocyte migration observed during recellularization.

GAGs are involved in different biological processes including extracellular matrix-cell interaction and activation of various chemokines [39]. They are stained blue on histology using Movat Pentachrome. Interestingly, from the 31^st^ day of recellularization, a blue coloration in the recellularized zones appeared on the Movat Pentachrome histological staining in 4 of 20 bronchi studied. These findings suggest the secretion of GAGs by the ASM cells, which are known to be secretory of these mucopolysaccharides [40].

No decellularization method is, to date, able to offer a complete elimination of the cellular material [41]. Thus, four criteria are used to assess the quality of a decellularized matrix: 1) maintenance of matrixial structural protein content, 2) absence of cellular material on histological staining, 3) double stranded DNA should be less than 50 ng/mg tissue and 4) less than 200 bp in length [42, 43]. In agreement with these reports, the bronchial extracellular matrix was histologically free of cellular material with a global maintenance of the bronchial architecture. The DNA concentrations we obtained by fluorometry and electrophoresis were below the thresholds recommended to avoid in vitro cytocompatibility problems [43]. However, in the context of regenerative medicine, Allman *et al.* showed that the immunoreactivity of the remnant DNA in a transplanted decellularized matrix could induce graft acceptance through a Th2 type response [44].

The decellularization protocol that was implemented herein was previously shown to allow a better preservation of the extracellular matrix than other methods [11, 45]. The total proteins fluorometric quantification revealed a significant concentration decrease of matrixial proteins in the decellularized bronchi compared to the native ones. This variation was expected as different intracellular proteins are eliminated during this process but the effects on the matrixial proteins seem to depend on the nature of the treated tissue [46, 47]. It has been shown that some cytoskeletal proteins, including α-SMA and SMMHC, could be detected in decellularized matrices [11, 47, 48] and correspond to the cellular residues observed by electron microscopy [48]. Our results are in agreement with these findings as on confocal microscopy, our decellularized matrices showed staining for α-SMA in some smooth muscle matrix areas. Moreover, the decellularized bronchi maintained their general architecture and protein composition almost unchanged in collagen I and IV, elastic fibers and fibronectin based on histological evaluation and semi-quantitative immunohistochemical scoring of native and decellularized bronchi. Those observations were in agreement with previous reports assessing the maintenance of these matrixial proteins [11, 14, 45, 48, 49]. Collagens and fibronectin are important for recellularization. Among other roles, fibronectin allow cell adhesion to the ECM and collagens are needed for their infiltration into it [47]. Laminin and other matrixial proteins also have cellular adhesion properties and may affect cell behavior and phenotype [50].

To conclude, we obtained a decellularized bronchial ECM that was successfully, while incompletely, recellularized with primary mature ASM cells over 41 days of culture. We described a preferential colonization of the smooth muscle ECM by these cells. Other investigations would be necessary to identify the factors and proteins that may be implicated in the ASM cell-specific recellularization observed.

## Acknowledgements

The authors thank Dr. Guy Beauchamp for the statistical analysis and the Respiratory Health Network of Quebec (RHN) for the tissue bank creation.

## Ethical approval

All applicable international, national, and/or institutional guidelines for the care and use of animals were followed. The experimental protocol was approved by the ethical committee of the University of Montreal number Rech-1578.

## supplemental Figures legends

**S1 Fig. Time chart of the recellularizations.** R is indicating recellularization at day 0 and S is referring to sampling on the timeline.

**S2 Fig. DNA electrophoresis for native and decellularized bronchi.** The abbreviation N.Br is for native bronchi and D.Br for decellularized ones.

## References

1. Gibson PG. What do non-eosinophilic asthma and airway remodelling tell us about persistent asthma? Thorax. 2007;62(12):1034–6.

2. Chetta A, Marangio E, Olivieri D. Inhaled steroids and airway remodelling in asthma. Acta Biomed. 2003;74(3):121–5.

3. Cukierman E, Pankov R, Stevens DR, Yamada KM. Taking cell-matrix adhesions to the third dimension. Science. 2001;294(5547):1708–12.

4. Baharvand H, Hashemi SM, Kazemi Ashtiani S, Farrokhi A. Differentiation of human embryonic stem cells into hepatocytes in 2D and 3D culture systems in vitro. Int J Dev Biol. 2006;50(7):645–52.

5. Htwe SS, Harrington H, Knox A, Rose F, Aylott J, Haycock JW, et al. Investigating NF-kappaB signaling in lung fibroblasts in 2D and 3D culture systems. Respir Res [Internet]. 2015 Dec 1 PMC4666055]; 16:[1–9 pp.]. Available from: https://www.ncbi.nlm.nih.gov/pubmed/26619903.

6. Duval K, Grover H, Han LH, Mou Y, Pegoraro AF, Fredberg J, et al. Modeling Physiological Events in 2D vs. 3D Cell Culture. Physiology (Bethesda). 2017;32(4):266–77.

7. Kapalczynska M, Kolenda T, Przybyla W, Zajaczkowska M, Teresiak A, Filas V, et al. 2D and 3D cell cultures - a comparison of different types of cancer cell cultures. Arch Med Sci. 2018;14(4):910–9.

8. Bullone M, Lavoie JP. The equine asthma model of airway remodeling: from a veterinary to a human perspective. Cell Tissue Res [Internet]. 2019 Nov 12. Available from: https://www.ncbi.nlm.nih.gov/pubmed/31713728.

9. Herszberg B, Ramos-Barbon D, Tamaoka M, Martin JG, Lavoie JP. Heaves, an asthma-like equine disease, involves airway smooth muscle remodeling. J Allergy Clin Immunol. 2006;118(2):382–8.

10. Leclere M, Lavoie-Lamoureux A, Joubert P, Relave F, Setlakwe EL, Beauchamp G, et al. Corticosteroids and antigen avoidance decrease airway smooth muscle mass in an equine asthma model. Am J Respir Cell Mol Biol. 2012;47(5):589–96.

11. Wagner DE, Bonenfant NR, Sokocevic D, DeSarno MJ, Borg ZD, Parsons CS, et al. Three-dimensional scaffolds of acellular human and porcine lungs for high throughput studies of lung disease and regeneration. Biomaterials. 2014;35(9):2664–79.

12. Vargas A, Peltier A, Dube J, Lefebvre-Lavoie J, Moulin V, Goulet F, et al. Evaluation of contractile phenotype in airway smooth muscle cells isolated from endobronchial biopsy and tissue specimens from horses. Am J Vet Res. 2017;78(3):359–70.

13. Russell HK, Jr. A modification of Movat’s pentachrome stain. Arch Pathol. 1972;94(2):187–91.

14. Baiguera S, Del Gaudio C, Jaus MO, Polizzi L, Gonfiotti A, Comin CE, et al. Long-term changes to in vitro preserved bioengineered human trachea and their implications for decellularized tissues. Biomaterials. 2012;33(14):3662–72.

15. Maghsoudlou P, Georgiades F, Tyraskis A, Totonelli G, Loukogeorgakis SP, Orlando G, et al. Preservation of micro-architecture and angiogenic potential in a pulmonary acellular matrix obtained using intermittent intra-tracheal flow of detergent enzymatic treatment. Biomaterials. 2013;34(28):6638–48.

16. Langhans SA. Three-Dimensional in Vitro Cell Culture Models in Drug Discovery and Drug Repositioning. Front Pharmacol [Internet]. 2018 PMC5787088]; 9:[6 p.]. Available from: https://www.ncbi.nlm.nih.gov/pubmed/29410625.

17. Yamada KM, Cukierman E. Modeling tissue morphogenesis and cancer in 3D. Cell. 2007;130(4):601–10.

18. Fisher SA, Tam RY, Fokina A, Mahmoodi MM, Distefano MD, Shoichet MS. Photo-immobilized EGF chemical gradients differentially impact breast cancer cell invasion and drug response in defined 3D hydrogels. Biomaterials. 2018;178:751–66.

19. Edmondson R, Broglie JJ, Adcock AF, Yang L. Three-dimensional cell culture systems and their applications in drug discovery and cell-based biosensors. Assay and drug development technologies. 2014;12(4):207–18.

20. Totonelli G, Maghsoudlou P, Garriboli M, Riegler J, Orlando G, Burns AJ, et al. A rat decellularized small bowel scaffold that preserves villus-crypt architecture for intestinal regeneration. Biomaterials. 2012;33(12):3401–10.

21. Taylor DA, Sampaio LC, Cabello R, Elgalad A, Parikh R, Wood RP, et al. Decellularization of Whole Human Heart Inside a Pressurized Pouch in an Inverted Orientation. J Vis Exp [Internet]. 2018 Nov 26; (141). Available from: https://www.ncbi.nlm.nih.gov/pubmed/30531712.

22. Xue A, Niu G, Chen Y, Li K, Xiao Z, Luan Y, et al. Recellularization of well-preserved decellularized kidney scaffold using adipose tissue-derived stem cells. J Biomed Mater Res A. 2018;106(3):805–14.

23. Stahl EC, Bonvillain RW, Skillen CD, Burger BL, Hara H, Lee W, et al. Evaluation of the host immune response to decellularized lung scaffolds derived from alpha-Gal knockout pigs in a non-human primate model. Biomaterials. 2018;187:93–104.

24. Crabbe A, Liu Y, Sarker SF, Bonenfant NR, Barrila J, Borg ZD, et al. Recellularization of decellularized lung scaffolds is enhanced by dynamic suspension culture. PLoS One [Internet]. 2015 PMC4427280]; 10(5):[e0126846 p.]. Available from: https://www.ncbi.nlm.nih.gov/pubmed/25962111.

25. Doi R, Tsuchiya T, Mitsutake N, Nishimura S, Matsuu-Matsuyama M, Nakazawa Y, et al. Transplantation of bioengineered rat lungs recellularized with endothelial and adipose-derived stromal cells. Sci Rep [Internet]. 2017 Aug 16 PMC5559597]; 7(1):[8447 p.]. Available from: https://www.ncbi.nlm.nih.gov/pubmed/28814761.

26. da Palma RK, Fratini P, Schiavo Matias GS, Cereta AD, Guimaraes LL, Anunciacao ARA, et al. Equine lung decellularization: a potential approach for in vitro modeling the role of the extracellular matrix in asthma. J Tissue Eng. 2018;9:1–11.

27. Burgstaller G, Sengupta A, Vierkotten S, Preissler G, Lindner M, Behr J, et al. Distinct niches within the extracellular matrix dictate fibroblast function in (cell free) 3D lung tissue cultures. Am J Physiol Lung Cell Mol Physiol. 2018;314(5):708–23.

28. McClure MJ, Cohen DJ, Ramey AN, Bivens CB, Mallu S, Isaacs JE, et al. Decellularized muscle supports new muscle fibers and improves function following volumetric injury. Tissue Eng Part A. 2018;24(15-16):1228–41.

29. Lyon CA, Koutsouki E, Aguilera CM, Blaschuk OW, George SJ. Inhibition of N-cadherin retards smooth muscle cell migration and intimal thickening via induction of apoptosis. J Vasc Surg. 2010;52(5):1301–9.

30. Wang X, Du C, He X, Deng X, He Y, Zhou X. MiR-4463 inhibits the migration of human aortic smooth muscle cells by AMOT. Biosci Rep [Internet]. 2018 Oct 31 PMC6147913]; 38(5). Available from: https://www.ncbi.nlm.nih.gov/pubmed/29752344.

31. Parameswaran K, Radford K, Zuo J, Janssen LJ, O’Byrne PM, Cox PG. Extracellular matrix regulates human airway smooth muscle cell migration. Eur Respir J. 2004;24(4):545–51.

32. Madison JM. Migration of airway smooth muscle cells. Am J Respir Cell Mol Biol. 2003;29(1):8–11.

33. Scarritt ME, Pashos NC, Motherwell JM, Eagle ZR, Burkett BJ, Gregory AN, et al. Re-endothelialization of rat lung scaffolds through passive, gravity-driven seeding of segment-specific pulmonary endothelial cells. J Tissue Eng Regen Med. 2018;12(2):786–806.

34. Uygun BE, Yarmush ML, Uygun K. Application of whole-organ tissue engineering in hepatology. Nat Rev Gastroenterol Hepatol. 2012;9(12):738–44.

35. La Rocca G, Anzalone R, Magno F, Corrao S, Carbone M, Loria L, et al. New perspectives on the roles of proteinases and lung structural cells in the pathogenesis of chronic obstructive pulmonary disease. In: Gerbino A, Zummo G, Crescimanno G, editors. Experimental medicine reviews Plumelia Ricerca ed 2007. p. 328.

36. Wachi H, Seyama Y, Yamashita S, Suganami H, Uemura Y, Okamoto K, et al. Stimulation of cell proliferation and autoregulation of elastin expression by elastin peptide VPGVG in cultured chick vascular smooth muscle cells. FEBS Lett. 1995;368(2):215–9.

37. Ribeiro SM, Poczatek M, Schultz-Cherry S, Villain M, Murphy-Ullrich JE. The activation sequence of thrombospondin-1 interacts with the latency-associated peptide to regulate activation of latent transforming growth factor-beta. The Journal of biological chemistry. 1999;274(19):13586–93.

38. Senior RM, Griffin GL, Mecham RP, Wrenn DS, Prasad KU, Urry DW. Val-Gly-Val-Ala-Pro-Gly, a repeating peptide in elastin, is chemotactic for fibroblasts and monocytes. J Cell Biol. 1984;99(3):870–4.

39. Papakonstantinou E, Karakiulakis G. The ‘sweet’ and ‘bitter’ involvement of glycosaminoglycans in lung diseases: pharmacotherapeutic relevance. Br J Pharmacol. 2009;157(7):1111–27.

40. Nigro J, Wang A, Mukhopadhyay D, Lauer M, Midura RJ, Sackstein R, et al. Regulation of heparan sulfate and chondroitin sulfate glycosaminoglycan biosynthesis by 4-fluoro-glucosamine in murine airway smooth muscle cells. The Journal of biological chemistry. 2009;284(25):16832–9.

41. Gilbert TW, Freund JM, Badylak SF. Quantification of DNA biologic scaffold materials. The journal of surgical research. 2009;1(152):135–9.

42. Gilpin A, Yang Y. Decellularization Strategies for Regenerative Medicine: From Processing Techniques to Applications. Biomed Res Int [Internet]. 2017 PMC5429943]; 2017:[9831534 p.]. Available from: https://www.ncbi.nlm.nih.gov/pubmed/28540307.

43. Crapo PM, Gilbert TW, Badylak SF. An overview of tissue and whole organ decellularization processes. Biomaterials. 2011;32(12):3233–43.

44. Allman AJ, McPherson TB, Badylak SF, Merrill LC, Kallakury B, Sheehan C, et al. Xenogeneic extracellular matrix grafts elicit a TH2-restricted immune response. Transplantation. 2001;71(11):1631–40.

45. Tsuchiya T, Sivarapatna A, Rocco K, Nanashima A, Nagayasu T, Niklason LE. Future prospects for tissue engineered lung transplantation: decellularization and recellularization-based whole lung regeneration. Organogenesis. 2014;10(2):196–207.

46. Grauss RW, Hazekamp MG, Oppenhuizen F, van Munsteren CJ, Gittenberger-de Groot AC, DeRuiter MC. Histological evaluation of decellularised porcine aortic valves: matrix changes due to different decellularisation methods. Eur J Cardiothorac Surg. 2005;27(4):566–71.

47. Gilbert TW, Sellaro TL, Badylak SF. Decellularization of tissues and organs. Biomaterials. 2006;27(19):3675–83.

48. Bonvillain RW, Danchuk S, Sullivan DE, Betancourt AM, Semon JA, Eagle ME, et al. A nonhuman primate model of lung regeneration: detergent-mediated decellularization and initial in vitro recellularization with mesenchymal stem cells. Tissue Eng Part A. 2012;18(23-24):2437–52.

49. Chani B, Puri V, Sobti RC, Jha V, Puri S. Decellularized scaffold of cryopreserved rat kidney retains its recellularization potential. PLos One [Internet]. 2017 PMC5340383]; 12(3). Available from: https://www.ncbi.nlm.nih.gov/pubmed/28267813.

50. Halper J, Kjaer M. Basic components of connective tissues and extracellular matrix: elastin, fibrillin, fibulins, fibrinogen, fibronectin, laminin, tenascins and thrombospondins. Adv Exp Med Biol. 2014;802:31–47.

