## Supplementary figures and images for "Recellularization of bronchial extracellular matrix with primary bronchial smooth muscle cells"

### Supplemental figure 1

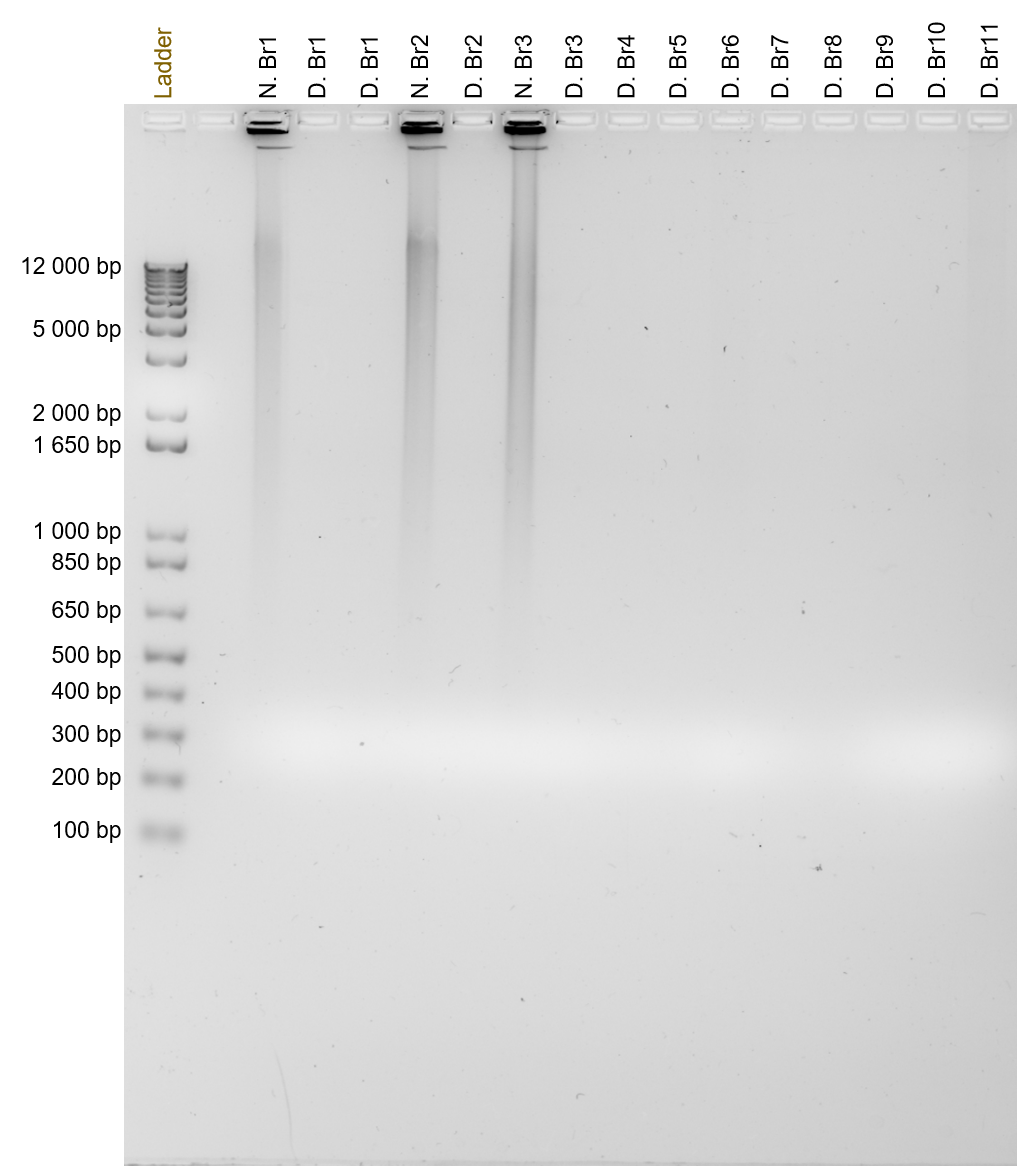

### Supplemental figure 2

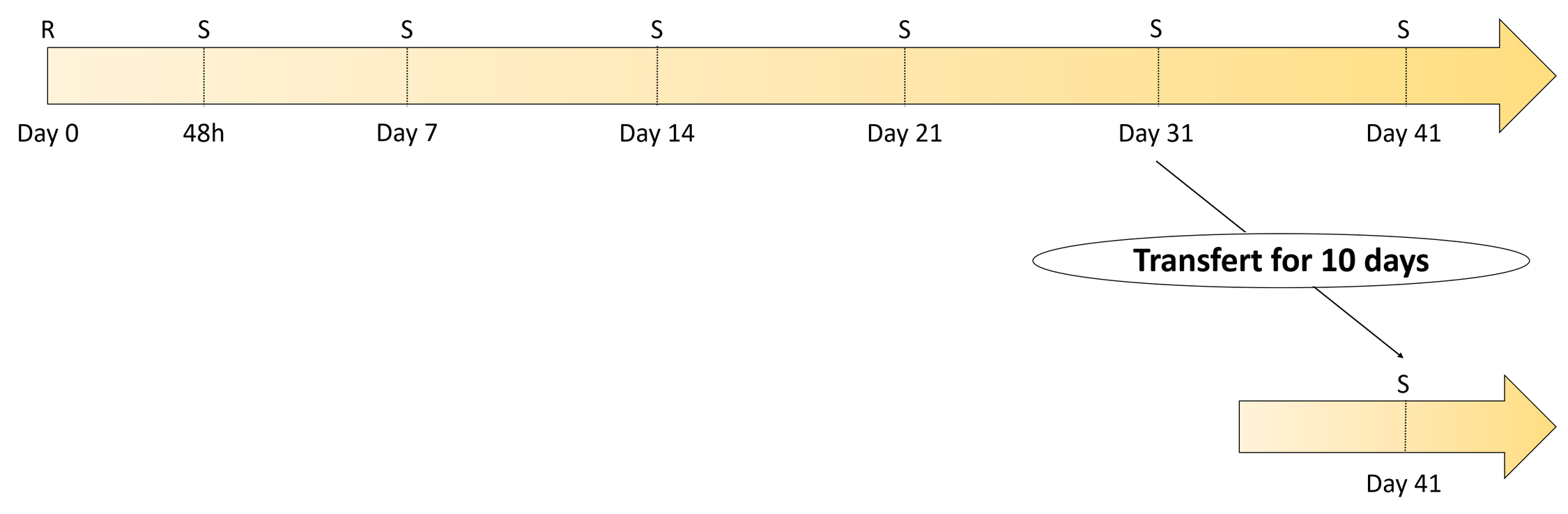
